# Glutamate, GABA, and dense-core vesicle secretion regulate predatory feeding in the nematode *Pristionchus pacificus*

**DOI:** 10.1101/2024.06.18.599456

**Authors:** Ageha Onodera, Andrew Zhang, Takahiro Chihara, Ralf J. Sommer, Misako Okumura

## Abstract

Nematodes are one of the most diverse groups of organisms, found in various environments, and exhibit various feeding behaviours. *Pristionchus pacificus* displays two types of feeding behaviours: predatory and bacterial feeding. Previously, we showed that the neurotransmitter serotonin plays important roles in regulating predatory feeding in *P. pacificus.* However, the role of other neurotransmitters in predatory feeding remains unclear. Using the CRISPR/Cas9 system, we generated mutants of the dense-core vesicle (DCV)-based neuropeptide secretion system and genes related to seven neurotransmitters. Predatory assays revealed that the vesicular glutamate transporter *Ppa-*EAT-4, glutamic acid decarboxylase *Ppa-*UNC-25 which is involved in GABA synthesis, and the calcium-dependent activator protein for DCV secretion *Ppa-*UNC-31, play roles in predatory feeding behaviour. We assessed the pharyngeal movement necessary for predation as well as locomotion rate in these mutants. While the *Ppa-eat-4* and *Ppa-unc-31* mutants decreased predation movement, the *Ppa-unc-25* mutant showed a reduction only in bacterial events compared to wild type animals. Additionally, *Ppa-unc-25* and *Ppa-unc-31* decreased in motor movement, potentially reducing predation efficiency. Together, these results suggest that glutamate, GABA and DCV secretion modulate feeding behaviours in *P. pacificus*. Our mutant collection of neurotransmitter-related genes will be useful for future analysis of neurobiology and behavioural evolution.

## Introduction

Animals have evolved a wide range of feeding behaviours to adapt to their ecology and diet. Nematodes are one of the most diverse groups of organisms, found in various environments and utilizing different food sources, including bacteria, fungi, plant, and other nematodes ^1,2^. The regulatory mechanisms of bacterial feeding behaviour have been studied in the model nematode *Caenorhabditis elegans*. In *C. elegans*, food bacteria are ingested by the pharynx, a neuromuscular organ that exhibits rhythmic muscle contractions known as pumping. The pharyngeal nervous system and combinations of neurotransmitters, such as serotonin, acetylcholine, glutamate, and GABA, and neuropeptides play roles in regulating pharyngeal pumping during bacterial feeding ^3,4^. For example, serotonin activates a cholinergic motor neuron, MC neuron, which stimulates pharyngeal pumping ^5–7^. Also, glutamate signalling modulates pharyngeal movement, especially in relaxation of pharyngeal muscles ^3,8^. Mutants of *Cel-unc-25* encoding the GABA synthesis enzyme exhibit a decrease in pumping rate during bacterial feeding, suggesting that GABA has a stimulatory role in pharyngeal pumping ^9^. Neuropeptides inhibit or promote pumping depending on conditions such as food availability in *C. elegans* ^9^. However, the mechanisms underlying the diverse feeding behaviours in other nematodes remain largely unknown.

The omnivorous nematode *Pristionchus pacificus*, which belongs to the family Diplogastridae, displays two distinct feeding behaviours: bacterial feeding and predatory feeding on other nematodes, with the latter representing an evolutionarily novel feeding behaviour ^10^. *P. pacificus* serves as a satellite model organism for comparison with *C. elegans*, facilitating integrative studies in evolutionary biology ^11–13^. Its annotated genome, partially understood synaptic connections, along with a variety of genetic tools such as transgenesis and reverse genetics with CRISPR/Cas9 ^14–19^, have advanced our understanding of the molecular mechanism of predation ^16,20–22^. Predatory feeding in *P. pacificus* depends on the presence of tooth-like denticles that allow it to penetrate the cuticle of its prey. The tooth-like denticles are part of a mouth-form dimorphism with two discrete forms ^23^. The eurystomatous (Eu) form is characterized by a wider buccal cavity and two tooth-like denticles, whereas the stenostomatous (St) form has a narrower buccal cavity and a single denticle. Correspondingly, each phenotype displays distinct feeding behaviours; Eu worms are able to kill prey with their movable teeth, while St worms cannot kill other worms but can only feed on dead corpses (Fig. 1A-C).

**Figure 1.**
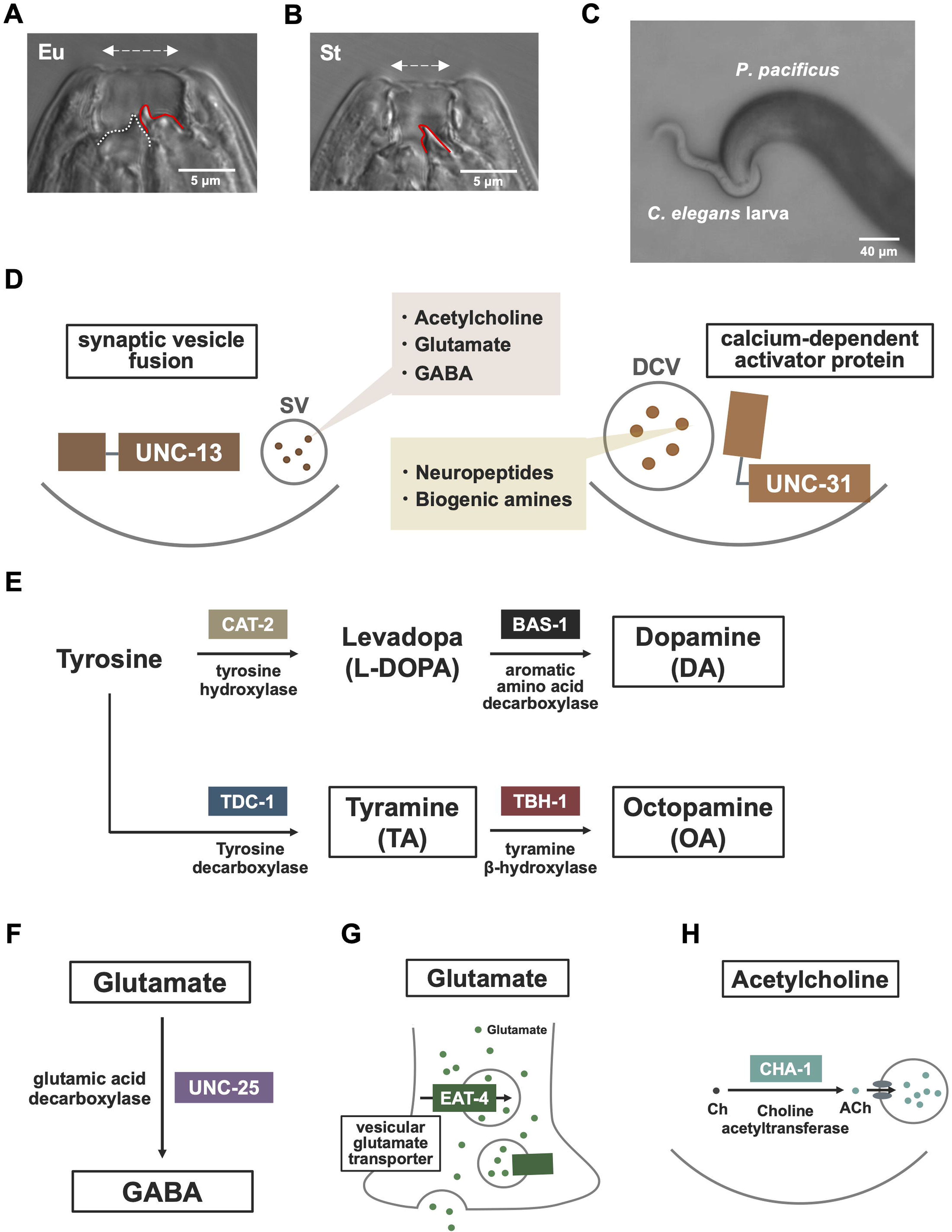
Mouth form plasticity and scheme of neurotransmitter functions. (A, B) DIC images of the mouth dimorphism in *P. pacificus.* The eurystomatous worm has a wide buccal cavity, a dorsal tooth (red line), and a subventral tooth (white dotted line). The stenostomatous worm has a narrow buccal cavity and a single dorsal tooth (white line), but no subventral tooth. Scale bars are 5 μm. (C) Predatory behaviour in Eu worm towards *C. elegans* prey. Scale bar is 40 μm. (D) The scheme of synaptic vesicle fusion protein (UNC-13) in neurotransmitter secretion and calcium-dependent activator protein (UNC-31). UNC-13 is required for neurotransmitter secretion via synaptic vesicles (SVs). UNC-31 is responsible for the secretion of neuromodulators enclosed within dense-core vesicles (DCVs). (E) The synthesis scheme for dopamine, tyramine, and octopamine. Dopamine is synthesized from tyrosine by two conserved enzymes called tyrosine hydroxylase (CAT-2) and aromatic amino acid decarboxylase (BAS-1). Tyramine is produced from tyrosine via an enzyme, tyrosine decarboxylase (TDC-1), while octopamine is synthesized from tyramine by an enzyme, tyramine β-hydroxylase (TBH-1). (F) The scheme of GABA synthesis. GABA is synthesized from glutamate by an enzyme, glutamic acid decarboxylase (UNC-25). (G) The scheme of EAT-4 function as a vesicle glutamate transporter, which is related to glutamate transduction. (H) Choline acetyltransferase (CHA-1) functions in acetylcholine processing.

Serotonin, a neurotransmitter, plays an important role in regulating predation by coordinating the rhythms of predatory feeding in *P. pacificus*. During predation, the tooth and the pharyngeal muscles exhibit rhythmic movements, which is mediated by serotonin and a specific subset of serotonin receptors ^16,20^. Treatment with serotonin but not with other biogenic amines such as dopamine, octopamine, or tyramine can induce tooth movement in the absence of prey ^10^. Interestingly, however, even mutants of serotonin synthase do not completely lose their predatory behaviour ^15^, suggesting that other neurotransmitters may contribute to predation, but the details have not been elucidated.

Here, we generated a collection of mutants related to neurotransmitters and dense-core vesicle secretion using the CRISPR/cas9 system and investigated the function of neurotransmitters including dopamine, octopamine, tyramine, glutamate, and GABA and neuropeptides in predatory feeding in *P. pacificus*. We found that a vesicular glutamate transporter (*Ppa-eat-4*), a glutamic acid decarboxylase (*Ppa-unc-25*), and a calcium-dependent activator protein for secretion (*Ppa-unc-31*) exhibited a reduction in predatory feeding behaviour compared to the wild-type animals.

## Material and Methods

### Strain

All strains of *P. pacificus* were incubated at 20℃on NGM plates with *Escherichia coli*, OP50. All experiments were performed using young adult hermaphrodites. All of the strains used in this study are listed in supplemental table 1.

### Generation of mutants using CRISPR/Cas9 genome editing

CRISPR mutants were generated as previously reported ^24–26^. A mixture of gRNA and Cas9 protein was injected into the gonad of hermaphrodites. F1 genotypes were examined by Sanger sequencing or high-resolution melting (HRM) analysis. The target sequences for gRNAs and primers are listed in supplemental table 2. All mutants except for the *Ppa-unc-31* mutant were backcrossed with the wild-type strain PS312 three times before the analyses. The *Ppa-unc-31* mutant could not be backcrossed because the mutant showed egg-laying defect and locomotion defect.

### Corpse assay

The corpse assay was conducted as previously described ^10,27^. Freshly starved *C. elegans* (N2) were collected with M9 buffer and filtered twice with 20 µm Nylon Net Filter (Millipore, NY2004700). After centrifugation, 2 µl of the larvae suspension was placed on NGM assay plates without *E. coli* OP50 and allowed to stand until the larvae were evenly dispersed across the plate. Five predators were added to each assay plate. After 2 h, the number of corpses was counted. The mouth morphology of the predators was examined under a light microscope and the number of corpses per Eu predator was calculated. The corpse assay was conducted in a blind manner.

### Bite assay

The bite assay was performed as previously described ^10,27^. Briefly, young larvae were collected as mentioned above. Predators were transferred to the assay plate with *C. elegans* larvae and were given a 15-30 min recovery period. After that, the numbers of biting, killing, and feeding events were counted using a light stereomicroscope for 10 min. The mouth morphology of the predators was examined under a light microscope and the data from the Eu animals were used for the analysis. The bite assay was conducted in a blind manner.

### Pumping and tooth movement

Pumping and tooth movement were counted as previously described ^10,27^. In short, worms were transferred to 35-mm NGM plates with OP50 or with *C. elegans* larvae. After 15 min recovery time, pharyngeal pumping and tooth movement were video recorded using a camera (ORCA-Fusion, C14440) for 15 sec in at least 10 animals under each condition. The mouth morphology of the predators was examined under a light microscope and the results of the Eu worms were used for quantification.

### Locomotion assay

The locomotion assay was conducted as previously described (Sawin et al. 2000). In brief, animals were washed twice in S basal buffer to remove bacteria and then transferred to assay plates without bacterial lawn. After 5 min of recovery, the number of body bends was counted for 20 seconds.

### Statistical analysis

The data were processed and analysed using GraphPad Prism (Version 10.2.2) and Microsoft Excel. Error bars represent the SEM, and descriptions of all statistical tests used as well as the meaning of symbols can be found in the figure legends.

### Material and Data Availability

All materials generated in this study, including worm strains, are available upon request. Any additional information required to reanalyze the data reported in this paper is available from the corresponding author upon request.

## Results

### Generation of neurotransmitter mutants using the CRISPR/Cas9 system

To examine whether neurotransmitters and neuromodulators regulate predatory feeding behaviour in *P. pacificus*, we firstly generated mutants of genes related to secretion of small synaptic vesicles (SVs) and dense core vesicles (DCVs) (Fig. 1D). In the process of neurotransmitter secretion, protein complexes are crucial in organizing the secretory system at the presynaptic zone ^28^. A synaptic protein, UNC*-*13, is involved in secretion via SVs that release fast-acting neurotransmitters including acetylcholine, glutamate, and GABA ^29^. A calcium-dependent activator protein for secretion (CAPS) protein, UNC-31, mediates secretion of DCVs that enclose peptides and biogenic amines ^30^. We could generate *Ppa-unc-31* (PPA06017, annotation with El_Paco_annotation_v3) homozygous mutants in *P. pacificus. Ppa-unc-31* mutants showed the bag of worm phenotype (Suppl.Fig.S1) and reduced locomotion (Fig. 4), as in the *C. elegans* studies^31^. However, we found that homozygous *Ppa-unc-13* (PPA35949, PPA03214, and ppa_stranded_DN31403_c0_g1_i5 are presumed to be different genes in El_Paco_annotation_v3, but the iso-seq data identified linked transcripts, suggesting that they are a single gene) mutants were lethal during development and could not be used for further analysis (Fig. 1D).

We also generated mutants in genes encoding for proteins that are involved in the synthesis or release of each neurotransmitter. We obtained homozygous frame-shift mutants for *Ppa-cat-2* (PPA44132), *Ppa-tdc-1* (PPA21225), *Ppa-tbh-1* (PPA23723), and *Ppa-unc-25* (PPA24743), whose orthologs are responsible for synthesizing dopamine, tyramine, octopamine, and GABA, respectively ^32–34^ (Fig. 1E, 1F). We also knocked out *Ppa-eat-4* (PPA15025), which is involved in vesicular glutamate transport, related to glutamate neurotransmission ^8^ (Fig. 1G). We tried to isolate homozygous mutants for *Ppa-cha-1* (PPA11636), which encodes an acetyltransferase involved in acetylcholine production ^35^ (Fig. 1H). However, the homozygous *Ppa-cha-1* mutants with a frame-shift mutation did not hatch. Thus, we have not performed further analysis for the *Ppa-cha-1* mutants. Together, we generated a mutant collection of neurotransmitters and DCV secretion, and mutants in *Ppa-unc-31, Ppa-cat-2*, *Ppa-tdc-1*, *Ppa-tbh-1*, *Ppa-unc-25*, and *Ppa-eat-4* were used for the further analyses.

### Predatory feeding behaviour was decreased in *Ppa-eat-4, Ppa-unc-25,* and *Ppa-unc-31* mutants

Using these mutants, we conducted two predatory assays: the corpse assay and bite assay (Figure 2A). In the corpse assay, five *P. pacificus* predators were placed on each assay plate with thousands of *C. elegans* L2 larvae, and the number of corpses was counted after two hours. The mouth morphology of the predators was examined and the number of corpses per Eu predator was calculated. While there was no significant difference in the number of corpses between the wild type and *Ppa-cat-2*, *Ppa-tdc-1*, and *Ppa-tbh-1* mutants, the *Ppa-eat-4*, *Ppa-unc-25*, and *Ppa-unc-31* mutant strains killed fewer prey than wild type animals. To examine the details of the predatory behaviour more closely, we conducted the bite assay, where the numbers of three predatory feeding aspects, biting, killing, and, feeding, were counted for 10 min. Consistent with the results of the corpse assay, the *Ppa-cat-2*, *Ppa-tdc-1*, and *Ppa-tbh-1* mutants showed similar numbers of biting, killing, and feeding behaviours as wild type animals (Figure 2B-2D). The *Ppa-eat-4* and *Ppa-unc-25* mutants exhibited a reduction in all predatory events (Figure 2E-2F). These results suggest that glutamate and GABA play roles in predatory feeding behaviour. Contrary to the finding of the corpse assay, the number of predatory events in the *Ppa-unc-31* mutant was comparable to those of the wild type (Figure 2G), which may be due to the different duration of the two predatory assays.

**Figure 2.**
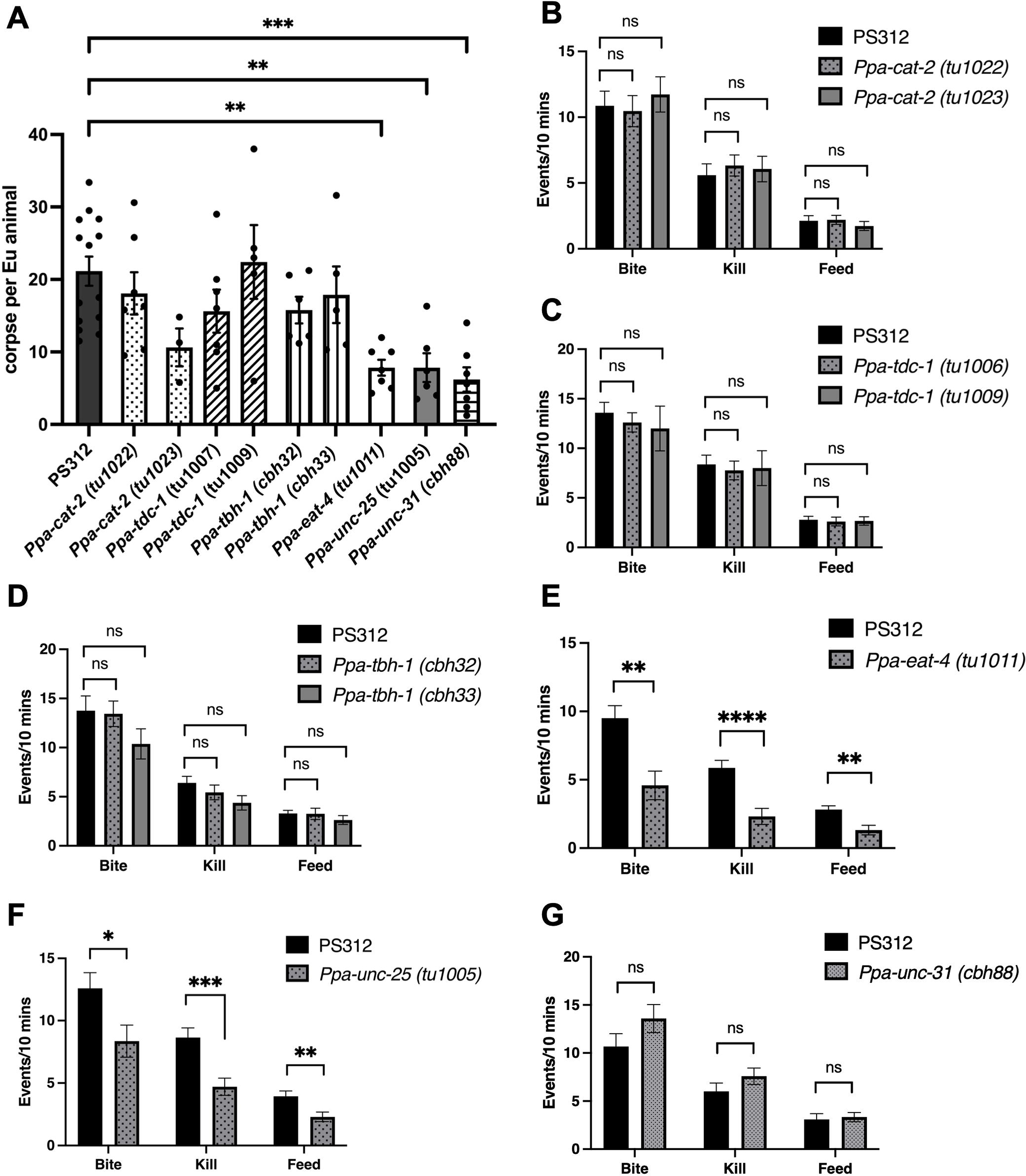
Two killing assays, corpse assay and biting assay revealed that glutamate and GABA play essential roles for predatory feeding behaviour. (A) Number of corpses of *C. elegans* larvae after 2 h incubations with five *P. pacificus* predators. PS312, *n* =14; *Ppa-cat-2* (tu1022), *n* = 6; *Ppa-cat-2* (tu1023), *n* = 3; *Ppa-tdc-1* (tu1007), *n* = 7; *Ppa-tdc-1* (tu1009), *n* = 5; *Ppa-tbh-1* (cbh32), n = 5; *Ppa-tbh-1* (cbh33), *n* = 5; *Ppa-eat-4* (tu1011), *n* = 7; *Ppa-unc-25* (tu1005), *n* = 6; *Ppa-unc-31* (cbh88), *n* = 8. Error bars are SEM. One-way ANOVA with Tukey’s multiple comparison tests. ns., not significant. **P<0.01. ***P<0.001 (B-G) Number of predation events in 10 min. PS312 *n* = 12-24; *Ppa-cat-2* (tu1022), *n* = 15; *Ppa-cat-2* (tu1023), *n* = 15; *Ppa-tdc-1* (tu1006), *n* = 13; *Ppa-tdc-1* (tu1009), *n* = 12; *Ppa-tbh-1* (cbh32), *n* = 16; *Ppa-tbh-1* (cbh33), *n* = 16; *Ppa-eat-4* (tu1011), *n* = 22; *Ppa-unc-25* (tu1005), n = 17; *Ppa-unc-31* (cbh88), n = 12. Error bars are SEM. One-way ANOVA with Tukey’s multiple comparison tests (B, C, D). Unpaired t test (E, F, G). ns., not significant. *P<0.05. **P<0.01. ***P<0.001

### *Ppa-eat-4* and *Ppa-unc-31* are involved in the modulation of tooth movement during predatory feeding

To clarify how these genes are involved in feeding behaviours, we first investigated the roles of these genes in regulating pharyngeal pumping rate in the presence of bacteria. Wild-type Eu animals exhibit higher pumping rate during bacterial feeding compared to that of predation ^10^. *Ppa-eat-4*, *Ppa-unc-25*, and *Ppa-unc-31* mutants showed a decrease in the pharyngeal pumping rate during bacterial feeding (Figure 3A). Given that in *C. elegans Cel-eat-4* and *Cel-unc-25* mutants exhibit a decrease in pumping rate during bacterial feeding ^9^, our results suggest conserved roles of glutamate and GABA in the modulation of the pumping rate during bacterial feeding. In contrast, the *Cel-unc-31* mutant did not show a reduction in pharyngeal pumping on food ^9^, whereas in *P. pacificus* the *Ppa-unc-31* mutant exhibited decreased pharyngeal pumping during bacterial feeding, suggesting that neuromodulators secreted by DCV have different roles between two species during bacterial feeding.

**Figure 3.**
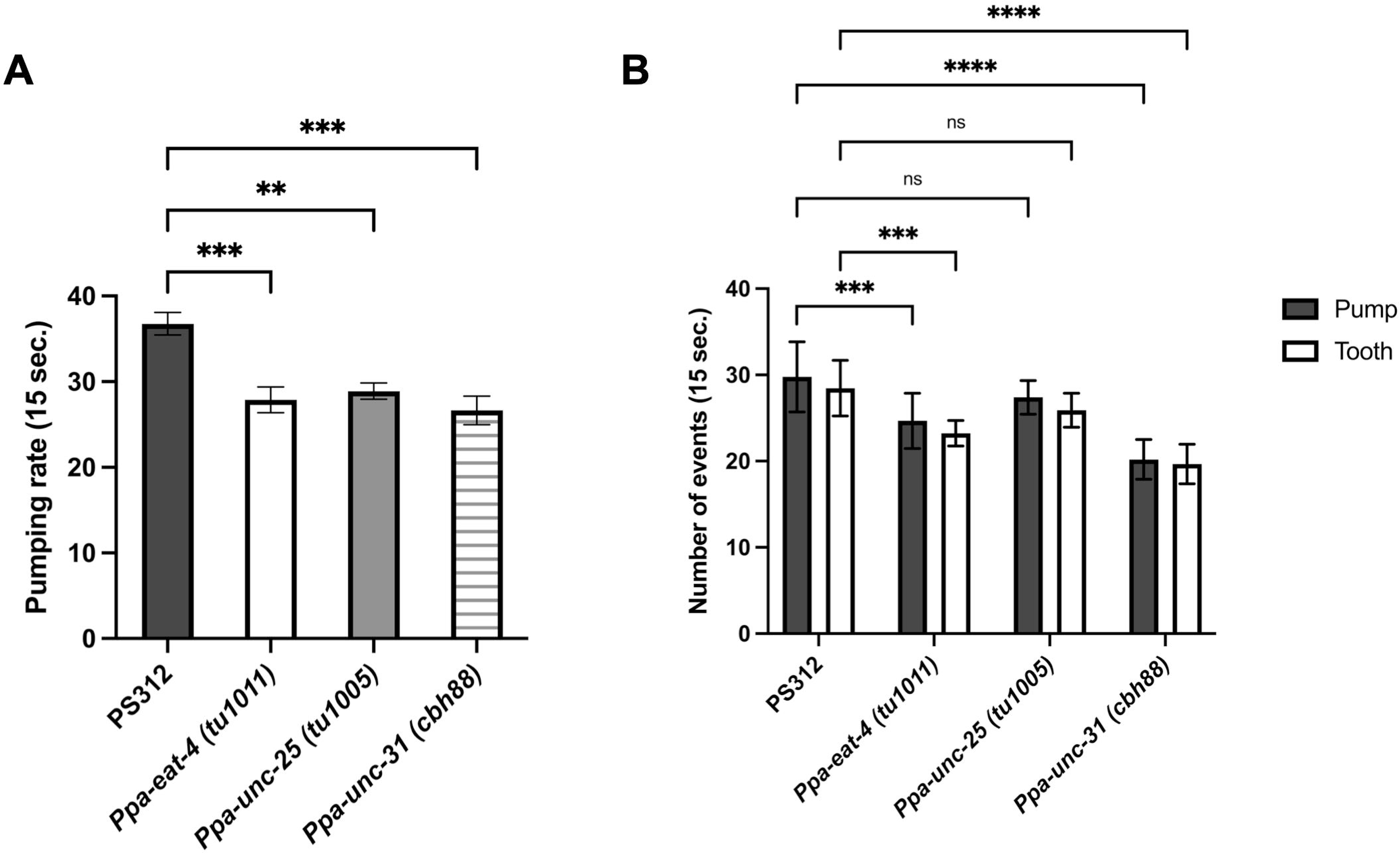
*Ppa-eat-4* and *Ppa-unc-31* mutants exhibited decreased pumping rate and tooth movement during predation. (A) Pharyngeal pumping rates of PS312, *Ppa-eat-4, Ppa-unc-25,* and *Ppa-unc-31* mutants on *E. coli* OP50. PS312, *n* = 10; *Ppa-eat-4* (tu1011), *n* = 14; *Ppa-unc-25* (tu1005), n = 10; *Ppa-unc-31* (cbh88), n = 8. Error bars are SEM. One-way ANOVA with Tukey’s multiple comparison tests. ns., not significant. **P<0.01. ***P<0.001. (B) Pumping rate and tooth movement during the predatory feeding behaviour. PS312, *n* = 13; *Ppa-eat-4* (tu1011), *n* = 9; *Ppa-unc-25* (tu1005), n = 10; *Ppa-unc-31* (cbh88), n = 10. Error bars are SEM. Two-way ANOVA with Tukey’s multiple comparison tests. ns., not significant. ***P<0.001. ****P<0.0001.

We also counted pharyngeal pumping and tooth movement during predation, which are important movements for efficient killing ^16^. In wild-type animals, pharyngeal pumping and tooth movement occur in an almost one-to-one ratio during predation ^10^(Figure 3B). The *Ppa-eat-4* and *Ppa-unc-31* mutants showed a reduction in both tooth movement and pumping, but the ratio of pumping and tooth movement was not disrupted (Figure 3B). The frequencies of pumping and tooth movement in the *Ppa-unc-25* mutant were comparable to those observed in wild type (Figure 3B). These results suggest that the glutamate and neuromodulators secreted via DCVs are involved in the modulation of tooth movement and pharyngeal pumping rate during predation.

### Locomotory rate decreases in *Ppa-unc-25* and *Ppa-unc-31* mutants

As the *Ppa-unc-25* mutant was not defective in tooth movement and pumping during predation, it is possible that the decrease of predation is caused by locomotion defect. We observed the locomotory rate by counting body bends for 20 seconds on an assay plate without bacteria. The locomotion rate of the *Ppa-unc-25* and *Ppa-unc-31* mutants were significantly lower compared to that of the wild-type strain (Figure 4). In contrast, the *Ppa-eat-4* mutant exhibited normal locomotion rate. The lower locomotory rate may potentially influence the efficiency of the predatory killing in the *Ppa-unc-25* and *Ppa-unc-31* mutants.

**Figure 4.**
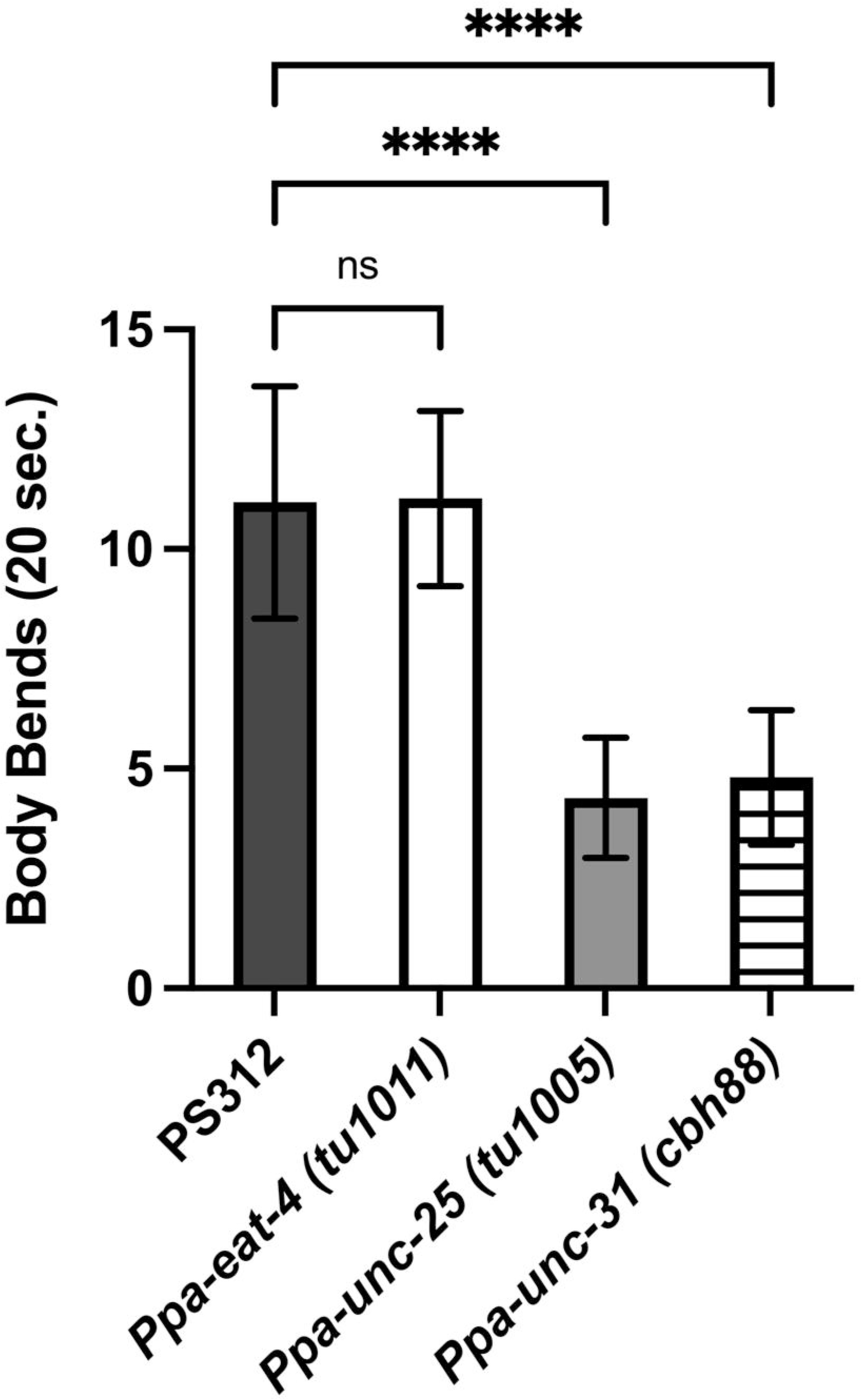
A decrease in locomotor activity was observed in *Ppa-unc-25* and *Ppa-unc-31* mutants. Number of body bends during 20 sec in PS312, *Ppa-eat-4, Ppa-unc-25,* and *Ppa-unc-31* mutants without OP50. PS312, *n* = 16; *Ppa-eat-4* (tu1011), *n* = 13; *Ppa-unc-25* (tu1005), n = 12; *Ppa-unc-31* (cbh88), n = 24. Error bars are SEM. One-way ANOVA with multiple comparison tests. ns., not significant. ****P<0.0001

**Figure 5.**
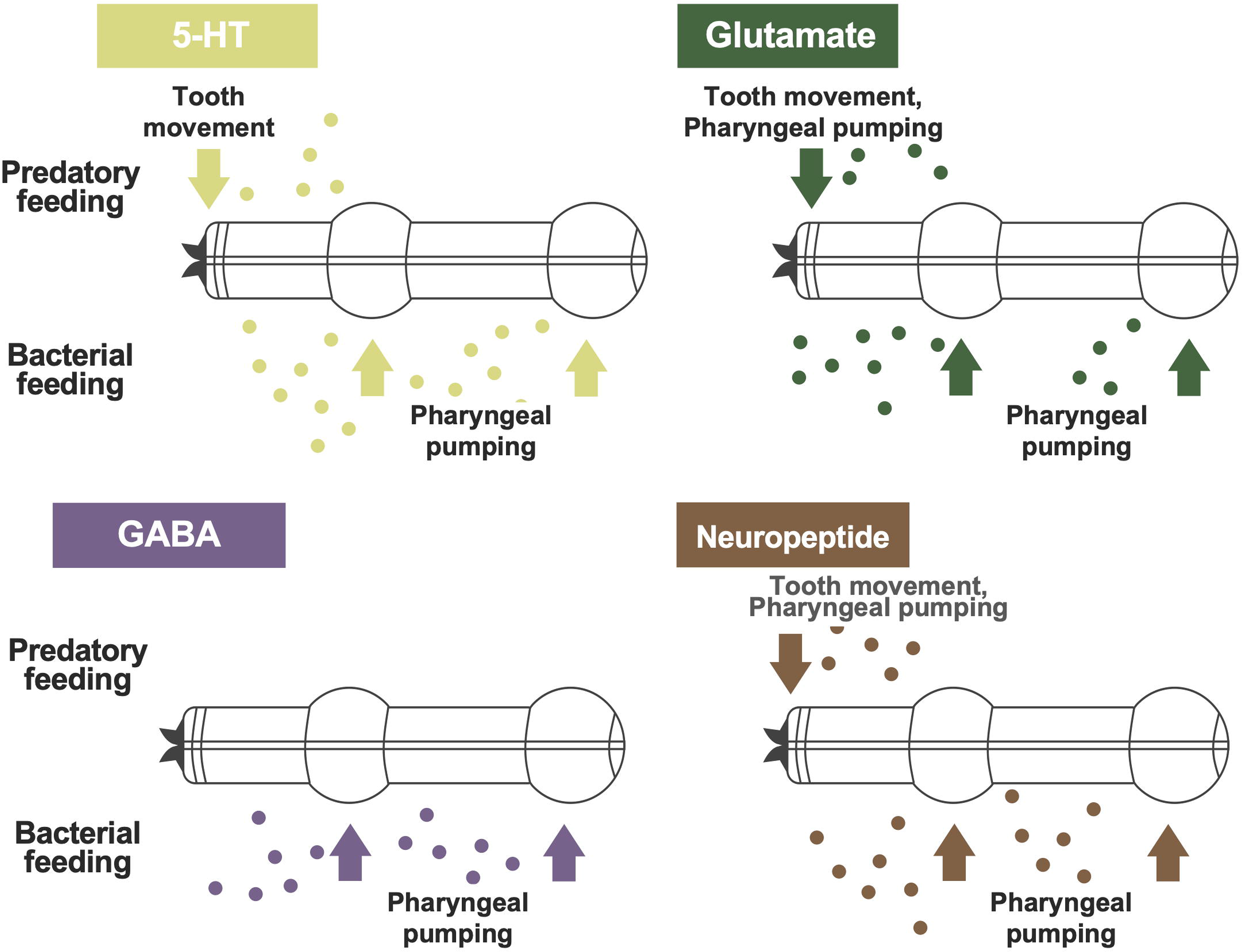
Schematic model of the neurotransmitter regulating system for the feeding behaviours in *P. pacificus*. A combination of neurotransmitters and neuromodulators contributes to both predatory and bacterial feeding behaviours. Previous studies revealed that serotonin (5-HT) plays a role in coordinating both tooth movement and pharyngeal muscle during predation, and stimulates pharyngeal pumping during bacterial feeding. Glutamate modulates tooth movement during predation and pharyngeal pumping on bacterial lawn. GABA is involved in pharyngeal muscle movement during bacterial feeding. Neuromodulators secreted via DVC have a role in both predatory feeding and bacterial feeding.

## Discussion

Nematodes exhibit a variety of feeding behaviours to adapt to the diversity of their habitat. In *C. elegans*, the combination of neurotransmitters and neuropeptides controls bacterial feeding behaviour in response to environmental conditions such as the presence of food or starvation ^9^. However, regulatory mechanisms of other feeding behaviours in other nematodes remain unclear. In this study, we generated a mutant collection for genes related to neurotransmitters and DCV secretion and examined their functions in predatory feeding in *P. pacificus*. The predatory assays revealed that glutamate, GABA, and neuromodulators secreted by DCVs play roles in predatory behaviour.

In *C. elegans Cel-eat-4* mutant, the frequency of pharyngeal pumping is reduced in the presence of food ^9^. M3 glutamatergic pharyngeal neurons activated by serotonin induce relaxation of the pharyngeal muscles ^3,8^. Although it is not known whether M3 neurons are glutamatergic in *P. pacificus*, M3 neurons make synapses with pm4 pharyngeal muscles in both species ^36^. These findings suggest that glutamate may have a common function in pharyngeal muscle regulation between the two species.

The *Ppa-unc-25* mutant showed a reduced frequency of predation, but the frequency of pumping and tooth movement during predation was comparable to that of the wild type. In *C. elegans*, GABA is important for locomotor behaviour ^37,38^. We observed a consistent phenotype in the *Ppa-unc-25* mutant in *P. pacificus*, implying that the reduced motility suppresses the successful predation in the *Ppa-unc-25* mutant. On the other hand, the frequency of pumping during bacterial feeding was reduced in *unc-25* mutants of both species. GABA expression has not been observed in the pharyngeal system of *C. elegans* ^39^, and it is unclear how GABA is involved in the regulation of pumping. Future studies, including GABA staining and identification and regulation of GABAergic neurons, should clarify how GABA is involved in the regulation of the two feeding behaviours in *P. pacificus*.

*Cel*-UNC-31 is involved in the secretion of neuromodulators, various neuropeptides and biogenic amines by DCV, and the *Ppa-unc-31* mutant also showed reduced pumping and tooth movement, and reduced locomotion, which may be involved in efficient predatory behaviour. Interestingly, the *Ppa-unc-31* mutant exhibited the predation defect in the corpse assay but not in the bite assay, which may be due to the difference in the duration between the two assays. In *C. elegan*s, *Cel-unc-31* modulates behavioural states: dwelling, roaming, and quiescence ^40^ and the *Cel-unc-31* mutant increased quiescent and dwelling time. It is likely that *Ppa-unc-31* regulates the behavioural states in *P. pacificus,* and the bite assay, which lasts only ten minutes, may not reflect its role in behavioural state modulation. Future analysis using long-time imaging of predatory behaviour and mutant analyses of neuropeptides would reveal whether the behavioural states modulated by neuropeptides influence predation.

We observed lethality in homozygous mutants in acetylcholine modification (*Ppa-cha-1*) and in a component of neurotransmitter secretion involved in syntaxin conformational change (*Ppa-unc-13*). In particular, we observed lethality after J2 moulting in *Ppa-unc-13* mutants, suggesting that neurotransmitter secretion via UNC-13 may be essential for larval development in *P. pacificus*. The auxin-inducible degradation (AID) system can rapidly degrade proteins in a variety of cell types in a spatial and temporal manner ^41,42^. The use of such a system in the future will advance the analysis of lethal genes, such as *Ppa-unc-13* and *Ppa-cha-1*.

Finally, our mutants could lead to the elucidation of the control mechanisms of not only predation but also other behaviours and the evolution of behaviour. For example, *P. pacificus* has a necromenic association with host beetles, and *P. pacificus* is primarily attracted by a beetle pheromone, which is different from C. elegans chemotaxis behaviour ^19,43^. It has also been reported that *C. elegans* and *P. pacificus* have different regulatory mechanisms even when they exhibit similar behaviours such as aggregation ^44,45^. Thus, our mutants may help to understand the neural regulation of the beetle-worm interaction and social behaviour, as well as other behaviours, which may promote our understanding of behavioural evolution.

## Acknowledgements

This work was supported by the Max Planck Society to R.J.S., JSPS Postdoctoral Fellowships for Research Abroad, JSPS KAKENHI (grant number 18K14716, 20K15903), AMED (grant number JP22gm6310003), and JST FOREST Program (grant number JPMJFR214M) to M.O., the Frontier Development Program for Genome Editing, the Astellas Foundation for Research on Metabolic Disorders, Naito Foundation, and JSPS KAKENHI (grant numbers 21H02479 and 21K18236) to T.C., and STEM Female Research Fellow in the science and technology fields at Hiroshima University to A. O. This work was conducted with the facility in the Natural Science Centre for Basic Research and Development (N-BARD) at Hiroshima University. We thank all the members of Sommer lab and Chihara lab for their kind support and discussion.

---

Supplemental Figure 1

*Ppa-unc-31* mutants exhibited the bag of worm phenotype

*Ppa-unc-31* mutants contain large numbers of larvae in their bodies.

Supplemental table 1

A list of the mutant strains generated in this study.

Supplemental table 1

A list of the gRNA target sequences and primers used in this study.

